# Adrenergic agonist induces rhythmic firing in quiescent cardiac preganglionic neurons in nucleus ambiguous via activation of intrinsic membrane excitability

**DOI:** 10.1101/276998

**Authors:** Isamu Aiba, Jeffrey L. Noebels

**Affiliations:** Department of Neurology, Baylor College of Medicine Houston TX 77030

## Abstract

Cholinergic vagal nerves projecting from neurons in the brainstem nucleus ambiguus (NAm) play a predominant role in cardiac parasympathetic pacemaking control. Central adrenergic signaling modulates the tone of this vagal output; however the exact excitability mechanisms are not fully understood. We investigated responses of NAm neurons to adrenergic agonists using *in vitro* mouse brainstem slices. Preganglionic NAm neurons were identified by Chat-tdtomato fluorescence in young adult transgenic mice and their cardiac projection confirmed by retrograde dye tracing. Juxtacellular recordings detected sparse or absent spontaneous action potentials (AP) in NAm neurons. However bath application of epinephrine or norepinephrine strongly and reversibly activated most NAm neurons regardless of their basal firing rate. Epinephrine was more potent than norepinephrine, and this activation largely depends on α1-adrenoceptors. Interestingly, adrenergic activation of NAm neurons does not require an ionotropic synaptic mechanism, since postsynaptic excitatory or inhibitory receptor blockade did not occlude the excitatory effect, and bath-applied adrenergic agonists did not alter excitatory or inhibitory synaptic transmission. Instead, adrenergic agonists significantly elevated intrinsic membrane excitability to facilitate generation of recurrent action potentials. T-type calcium current (ICaT) and hyperpolarization-activated current (Ih) are involved in this excitation pattern, while not required for spontaneous AP induction by epinephrine. In contrast, pharmacological blockade of persistent sodium current (INaP) significantly inhibited the adrenergic effects. Our results demonstrate that central adrenergic signaling enhances the intrinsic excitability of NAm neurons, and persistent sodium current is required for this effect. This central balancing mechanism may counteract excessive peripheral cardiac excitation during increased sympathetic tone.

**New & Noteworthy:** Cardiac preganglionic cholinergic neurons in the Nucleus ambiguus (NAm) are responsible for slowing cardiac pacemaking. This study identified that adrenergic agonists can induce rhythmic action potentials in otherwise quiescent cholinergic NAm preganglionic neurons in brainstem slice preparation. The modulatory influence of adrenaline on central parasympathetic outflow may contribute to both physiological and deleterious cardiovascular regulation.

## Introduction

Central parasympathetic autonomic regulation is predominantly mediated by vagal preganglionic fibers originating in the ventrolateral brainstem medulla oblongata. The caudal portion of the nucleus ambiguus (NAm) contributes the majority of axons in the cardiac vagus nerve and is responsible for cholinergic reduction of heart rate. Chronically reduced vagal tone is implicated in an increased risk of ventricular tachycardia and fibrillation, while enhanced tone has antiarrhythmic properties (Kalla et al. 2016). On the other hand, acute vagal hyperactivity may lead to bradycardia, asystoles, atrial fibrillation, vagal reflex syncope and, in severe cases, cardiac arrest (Alboni et al. 2011; Colman et al. 2004). Strong autonomic co-activation is also arrhythmogenic and evokes complex sympathetic-and parasympathetic-driven cardiac arrhythmias (Koizumi and Kollai 1981), which can be life-threatening in people with heart disease (Shattock and Tipton 2012).

Central adrenergic signaling is likely involved at several levels in the homeostatic response to stress autonomic regulation (Kvetnansky et al. 2009). The descending adrenergic neurons within the rostral ventrolateral medulla brainstem (e.g. A5, C1r groups) project to preganglionic cardiac sympathetic neurons in the intermediolateral cell column of the thoracic spinal cord to modulate cardiac sympathetic outflow (Loewy et al. 1979), and subsets of these adrenergic neurons may also project collaterals to the NAm (Byrum and Guyenet 1987; Kalia et al. 1985; Stocker et al. 1997) to modulate vagal outflow (Abbott et al. 2013). Adrenergic neurons in the locus coeruleus (A6 group) neurons also modulate local inhibitory synaptic transmission in the NAm (Wang et al. 2014). Given the need to maintain tight coordination for optimal sympathetic and parasympathetic balance, central adrenergic feedforward signaling may play important roles in determining cardiac autonomic regulation. However the mechanisms underlying adrenergic signaling directly onto preganglionic neurons in the NAm is not fully understood. Cardiac vagal neurons do not show pacemaking activity *in vitro* (Mendelowitz 1996); their activity is considered to be regulated by synaptic activation of postsynaptic ionotropic glutamate, GABA and glycine signaling, and this input is influenced by various neuromodulators (Dergacheva et al. 2009; Dergacheva et al. 2016; Dyavanapalli et al. 2013; Jameson et al. 2008; Philbin et al. 2010; Sharp et al. 2014). On the other hand, the intrinsic membrane excitability of NAm neurons is enhanced by non-synaptic neuromodulators, for example, aldosterone depolarization in neonatal rat neurons (Brailoiu et al. 2013b). Currently, the relative importance of synaptic versus intrinsic membrane excitability mechanisms on the discharge patterns of NAm neurons is not fully understood, nor is it known whether the mechanisms largely studied in immature animals are conserved in the young adult (Kasparov and Paton 1997).

The present *in vitro* study of young adult NAm cholinergic neurons in mouse brainstem slices shows that adrenergic agonists exert potent non-synaptic excitatory effects on preganglionic vagal neurons. While a majority of NAm neurons are quiescent in the absence of synaptic transmission, we found that adrenergic agonists can induce pacemaking activity. This adrenergic excitatory effect produced a robust rhythmic discharge pattern that did not depend upon ionotropic synaptic currents, but rather potentiated intrinsic membrane excitability. This mechanism may play a role in coordinate central regulation of sympathetic and parasympathetic autonomic system, and may be a useful target for treatment of abnormal cardiac conditions.

## Material and methods

### Animals

All experiments were conducted under the protocol approved by the IACUC at Baylor College of Medicine. Chat-IRES-cre mouse (JAX: stock No: 006410) and cre dependent tdtomato reporter mouse (Ai9, JAX: Stock No: 007909) were obtained from Jackson laboratory and crossed to generate Chat-cre (+/cre), tdtomato (+/flox or flox/flox) mouse line. Mice aged between P20-50 of both sexes were used.

### Retrograde labeling of cardiac premotor neurons in brainstem

Mice (Chat-cre, tdtomato) were deeply anesthetized with avertin (Tribromoethanol, 200 mg/kg i.p.) or 2% isoflurane. Skin overlying the precordial region was depiliated, cleansed, and a small vertical skin incision made along the sternal line. The thoracic wall was exposed and DiO suspension (30 mg/ml, 100 µl, 30% DMSO in saline) was slowly injected into the pericardial space via the intercostal spaces of the left 3-5^th^ ribs (~1.5 mm depth). Breathing pattern was carefully monitored to ensure absence of pneumothorax. The skin incision was sutured and the mouse allowed to recover for at least 1 week. For histological analysis, mice were cardiac-perfused with ice cold PBS followed by 4% PFA. Brain was extracted and kept in 4% PFA for overnight at 4°C, followed by further incubation in 30% sucrose until brain sinks. Brain was embedded in OCT compound and cut in 50-70 µm coronal sections with a cryostat. Brain sections were rinsed with PBS and mounted on glass slides. Fluorescence images were acquired by fluorescence microscopy (Nikon TE2000S) with the NIS element program and analyzed with ImageJ.

### In vitro Electrophysiology

Mice were deeply anesthetized with avertin (Tribromoethanol, 200 mg/kg i.p.), cardiac perfused with a brain cutting solution (in mM: 110 NMDG, 6 MgSO_4_, 25 NaHCO_3_, 1.25 Na_2_HPO_4_, 0.1 CaCl_2_ 3 KCl, 10 glucose, 0.4 ascorbate, 1 thiourea, saturated with 95% O_2_/5% CO_2_) and decapitated. The brain was quickly extracted in ice-cold cutting solution, the isolated hindbrain was glued to a mold, the cerebellum removed, and 200 µm thick coronal medulla slices were cut with a vibratome (Leica VT-1200). Slices were further hemisected at the midline, incubated in ACSF (in mM: 130 NaCl, 3 KCl, 1 MgSO_4,_ 2 CaCl_2_ 25 NaHCO_3_, 1.25 Na_2_HPO_4_, 10 glucose, 0.4 ascorbate, saturated with 95% O_2_/5% CO_2_) at 32°C for 1 hour, then kept in oxygenated ACSF at room temperature.

Recordings were made in a submerged chamber (RC27, Warner Instruments) continuously perfused with ACSF at 2.5 ml/min and 32-33°C. NAm was visually identified in the transparent regions located in the ventral medulla, and the cholinergic neurons were confirmed by presence of tdtomato fluorescence. All electrophysiological signals were amplified with a Multiclamp 200B amplifier, digitized and acquired with Clampex software (Molecular Devices). Data were analyzed with pClamp9 and Mini Analysis Program (Synaptosoft).

Cell attached patch recordings were made by voltage clamp recording with micro pipettes (1-4 MΩ) containing ACSF. Seal resistance was between 5-50 MΩ. Data were >1 Hz high pass filtered. In order to minimize perturbation of membrane potentials of recorded neurons, the command voltage was adjusted to a voltage at which holding current was within ±300 pA (Perkins 2006). Recordings were included for analysis only when neurons showed spontaneous action potentials, or silent at rest but with action potentials that could be evoked by bipolar stimulation (electrodes ~300 µm apart) of adjacent tissue. The maximum AP frequency calculated from multiple 10 seconds bins of recorded traces. The peak firing period typically lasts longer than 1 minute and the variability in the duration of induced AP firing does not critically affect the conclusion.

In whole cell recordings, the membrane resistance, capacitance and access resistance were determined using the membrane test protocol of Clampex software (10 mV, 20 ms rectangular test pulse, at -70 mV holding potential). The measured resistance primarily reflects leak current near the resting potential. In some recordings, data were <100 Hz low pass filtered. Cells with an access resistance <20 MΩ were accepted for analysis.

Current clamp recordings were made with potassium gluconate internal solution (in mM: 135 potassium gluconate, 10 HEPES, 1 MgCl_2_, 8 NaCl, 0.05 EGTA, 2 Mg-ATP, 0.3 Na-GTP, pH adjusted to 7.2 with KOH). Intrinsic membrane excitability was tested after adjustment of bridge balance. Responses to rectangular test pulses (-200 to +450 pA, 50 pA increment, 500 ms) were measured at resting potential or -70mV to avoid generation of spontaneous action potentials. Liquid junction potentials were adjusted by 10mV.

Voltage-clamp recordings were made with a cesium gluconate internal solution (in mM: 130 gluconic acid, 10 HEPES, 1 MgCl_2_, 8 NaCl, 10 BAPTA, 5 TEA, 5 QX314, 2 Mg-ATP,0.3 Na-GTP, pH adjusted to 7.2 with CsOH). Liquid junction potentials were adjusted by 10 mV. Low voltage-activated currents were evoked by the following current protocol: prepulse at -110 mv for 2 seconds followed by voltage step to various potentials (-70 to - 30mV with 10mV increment). When testing steady state inactivation (SST), T-type calcium current was evoked with hyperpolarizing prepulses (-110 to -50 mV with 10 mV increments) at 2 second intervals, followed by an activation voltage step to -50 mV. Both activation and SST I-V curves were fitted to a sigmoidal curve and V_50_ values were calculated. Maximum current amplitude was obtained from the activation curves. Persistent sodium current was recorded with a cesium gluconate internal solution without sodium channel blocker QX314. Cadmium (100 µM) was included in the superfusate to reduce voltage-gated calcium currents. Neurons were clamped at -80mV and slowly depolarized by a voltage ramp (20 mV/s) until +20 mV. After obtaining basal current, recordings were repeated in the presence of tetrodotoxin (TTX: 10 nM, 1 µM) or 30 µM riluzole to isolate the INaP component. Traces averaged from 3-6 sweeps were used for analysis.

### Drugs

Epinephrine, norepinephrine, phenylephrine, doxazosin, isoproterenol were from Sigma Aldrich USA. NBQX, gabazine (SR 95531), carveninolol and riluzole were obtained from Tocris. TTPA2 and TTX were purchased from Alomone Labs. Riluzole was dissolved in DMSO at 300 mM and stored at -20°C. Other drugs were directly dissolved in ACSF on the day of experiments. The selective T-type calcium channel blocker Z944 was kindly provided by Dr Terry Snutch (Tringham et al. 2012). GS967 was obtained from MedChemExpress.

### Experimental Design and Statistical Analysis

Both male and female mice were used. Because preliminary experiments did not detect a sex difference, data were pooled. All data are presented as mean ±standard deviation unless specifically mentioned.

Statistical analyses were conducted by Graphpad Prism and R software. Spontaneous AP frequency change between baseline and Epi exposure was tested by Wilcoxon matched-pairs signed rank test. For repetitive experiments (Figure 3, 7 and 9), single neurons were exposed to Epi twice, first in control solution and second in the presence of test compound. Quantitative statistical significance in repeated measurements was tested by repeated-measures ANOVA with post hoc Tukey’s multiple comparisons test. Result of ANOVA test was provided by F statistic with p-values in figure legends. In some pharmacological analyses (Figure 3, 4, 7, 8), the fraction of cells that did not respond to epinephrine in the presence of drug treatment was tested by contingency table analysis with Fisher’s exact test. Cells responding by a >10% increase in their AP frequency were considered to be activated by epinephrine.

## Results

### Putative cardiac premotor neurons in the nucleus ambiguus (NAm)

Preganglionic neurons in the NAm were identified using a transgenic mouse expressing tdtomato fluorescent protein under the control of the *choline acetyl transferase* (Chat) promoter (Chat-tdtomato). Their cardiac projection was verified by retrograde labeling with a tracer dye (DiO) injected into the pericardial sac (see methods) Figure 1A. All retrogradely labeled DiO^+^ cells were tdtomato positive and had a large soma diameter (>40 µm), while ~10% of the Chat-tdtomato^+^ cells were not labeled with retrograde tracers. These non-cardiac labeled cells typically had smaller somata and were sparsely distributed within and around the NAm (Figure 1B, n=3 mice). No neurons within the dorsomotor vagal nucleus (DMV) were retrograde-labeled. The lack of DMV labeling may indicate failure to label a subpopulation of cardiac projecting neurons with our method, but also indicates little misidentification of gastric preganglionic neurons within NAm (Grkovic et al. 2005).

**Figure 1.**
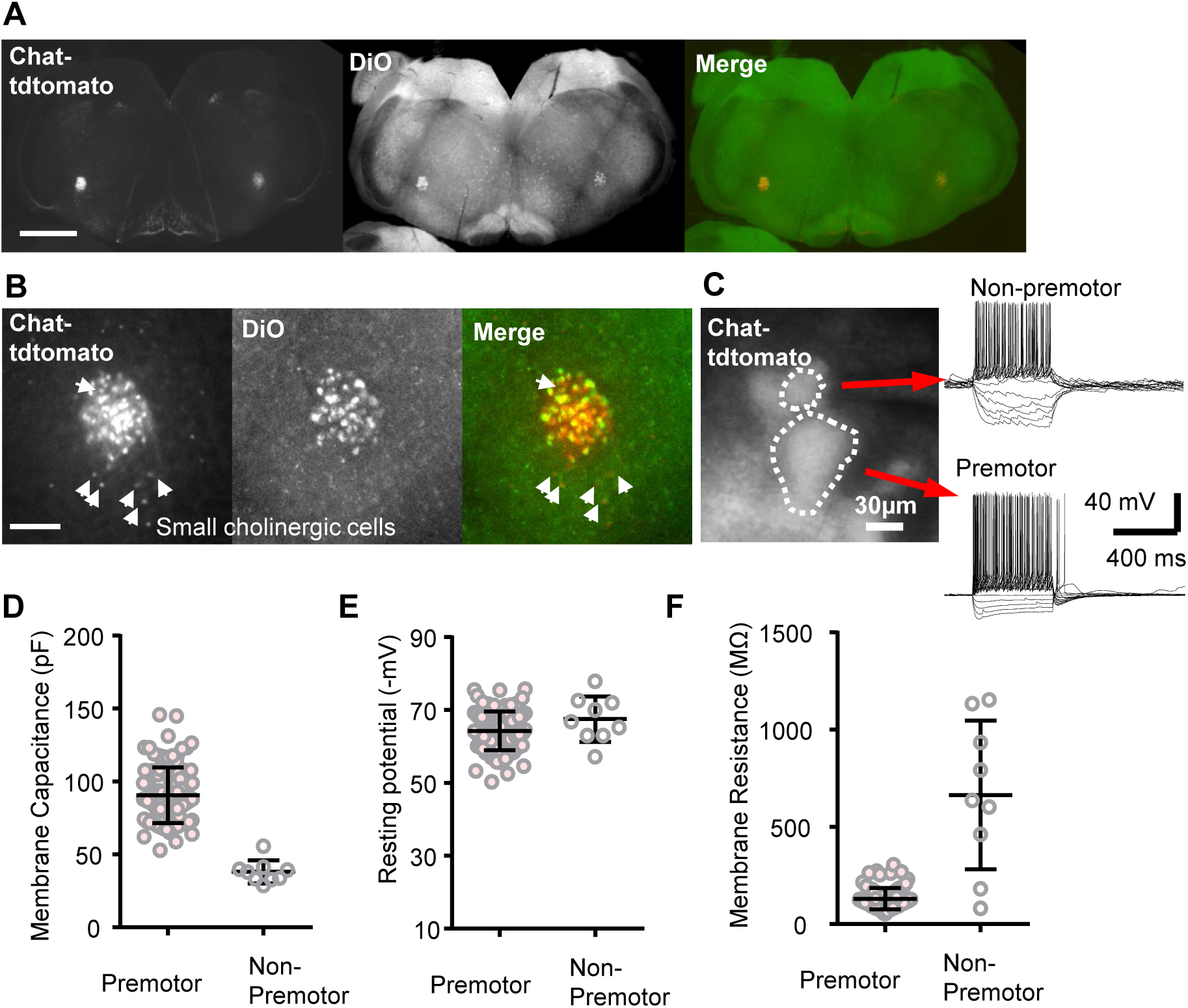
Chat-tdtomato+ labeling of preganglionic neurons within the Nucleus Ambiguus (NAm). **A&B** Retrograde tracer DiO was injected into the pericardiac sac, and cells colabeled with retrograde dye and Chat-tdtomato fluorescence were analyzed. Low magnification (4x, scale bar: 1 mm, **A**) and higher magnification (20x, scale bar: 400 µm, **B**) images. DiO was detected mostly in cells with a large soma and was absent in the majority of small Chat-tdtomato^+^ cells (arrow heads), suggesting that most preganglionic cells have a larger soma size. **C.** Unlabeled preganglionic and non-preganglionic Chat-tdtomato^+^ cells (threshold soma size 40 µm) could be readily distinguished in acute brainstem slices prepared from young adult animals. **D-F**. Preganglionic neurons (n=100, soma diameter >40 µm) and non-preganglionic (n=9) neurons showed distinct membrane excitability.

The electrophysiology of Chat-tdtomato^+^ cells was characterized *in vitro* using acute slices prepared from young adult (P20-P50) mouse brainstem. The two populations with different cell sizes could be also distinguished *in vitro* and showed distinct electrophysiological properties. Consistent with the difference in soma size, these two populations had different mean membrane capacitances (90.9±19.4 pF vs 36.6±4.4 pF, n=100, 9, Figure 1D). The resting potentials were similar (-64.5±5.4 mV vs -64.1±1.5 mV, Figure 1E). The membrane resistance of neurons with larger soma diameter was low and relatively homogenous, while membrane resistances of the smaller neurons differ across a larger range (130.0±53.8 MΩ vs 663.4±382.0 MΩ, Figure 1F). These results suggest that the Chat-tdtomato^+^ cells with a small soma are a mixed population and may be cholinergic local circuit or oropharynx/larynx projection neurons (Irnaten et al. 2001). Based on histology and electrophysiological characterizations, the Chat-tdtomato^+^ cells with larger soma (>40 µm diameter) are presumed to be the long range cardiac preganglionic neurons, and in subsequent experiments, all analyses were made from this cholinergic neuron group.

### Adrenergic agonists activate spontaneous rhythmic AP firing in NAm neurons in part via 1 and receptor mechanisms

Adrenergic signaling could regulate NAm cholinergic output through modulation of excitatory and inhibitory synaptic transmission, as shown by subtype selective adrenoceptor agonists/antagonists *in vitro* studies (Bateman et al. 2012; Boychuk et al. 2011; Philbin et al. 2010; Sharp et al. 2014). However the possibility of direct postsynaptic effects has not yet been clearly demonstrated. We recorded NAm AP firing using cell-attached microelectrode recordings. Under basal conditions, NAm cells showed a variety of spontaneous AP firing patterns in vitro, ranging from complete silence to active firing (2.07±2.43 Hz, n=111 neurons, Figure 2C). Bath application of epinephrine (hereafter Epi; 10 µM) or norepinephrine (hereafter NE; 10 µM) significantly enhanced spontaneous AP firing in NAm neurons (Figure 2A). Epinephrine increased the spontaneous action potential rate in a majority of these cells (92%; 102/111 cells), while the remainder (8%; 9/111 cells) displayed a ‘burst suppression’ firing pattern (Figure 2B). Due to the low chance of encountering the burst cells, further characterization focused on the regular-spiking cells. Linear regression analysis detected a positive correlation between firing rate in baseline and after Epi exposures across the regular spiking neurons (p=0.0002, Figure 2E).

**Figure 2.**
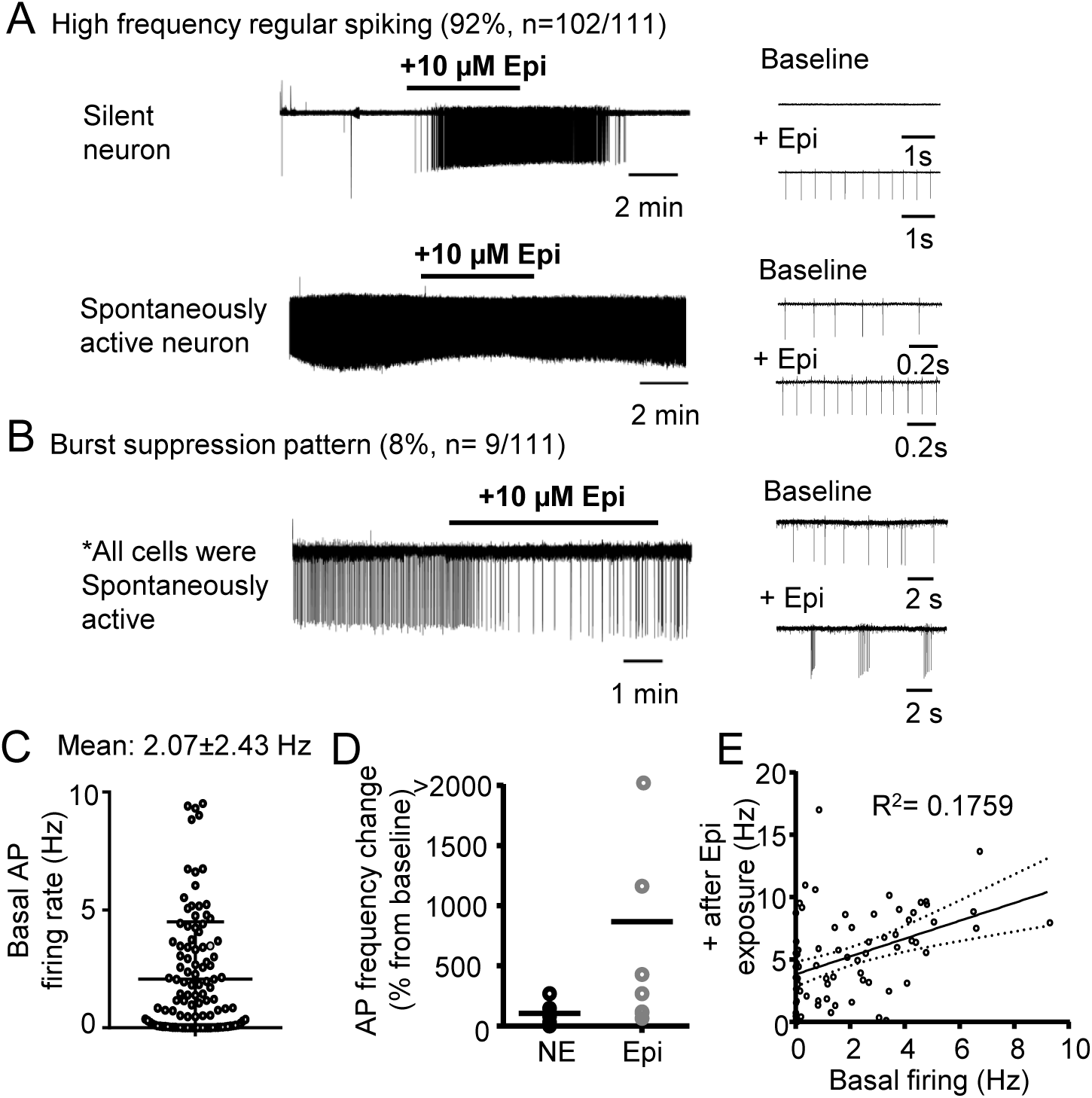
Adrenergic agonist activation of NAm neurons *in vitro.* **A.** Bath application of epinephrine (10 µM) reversibly increased spontaneous action potential rate in silent (top) and spontaneous active (bottom) neurons. **B.** In some NAm neurons (8%, 9/111 cells), epinephrine modulated NAm firing in a burst suppression pattern. All of the burst neurons were spontaneously active. **C.** Summary data of basal spontaneous AP rate of recorded neurons (n=111). **D.** Comparison of AP frequency changes by repetitive Epi and NE exposures. Norepinephrine had a less potent excitatory effect (n=7 neurons, repeated-measures ANOVA F(1, 12) = 19.62, p= 0.0008). **E.** A linear regression analysis of AP firing rate in baseline and after Epi exposure. A correlation was detected. R^2^ = 0.1759, p=0.0002. Solid line: regression line, dashed line: 95% confidential interval.

Epi had a stronger excitatory effect than NE. Comparison of efficacy was made by sequentially exposing single neurons with Epi and NE (4 neurons first exposed to Epi washout and then to NE, other 3 exposed to NE, washed, then to Epi). Epi increased spontaneous AP rate 9.7±1.4-fold, while NE increased AP 2.1±0.9-fold in the same neurons (n=7, Figure 2D). Due to its higher efficacy, the majority of subsequent experiments were performed using epinephrine as an agonist.

We next examined the adrenoceptors involved in adrenergic activation of NAm neurons. In order to ensure regular-spiking response to Epi, NAm neurons were first exposed to Epi, washed, and then re-exposed to Epi with or without a selective antagonist for each test compound.

In time controlled experiments, 1^st^ and 2^nd^ Epi exposures increased spontaneous AP rate of the NAm neurons to a similar degree (n=9, Figure 3A). In experiments with the pan-βreceptor antagonist propranolol (10 µM), wash-in of the antagonist reduced the basal spontaneous AP firing rate (76.3±27.8% reduction). The antagonist also blocked AP increases in 38% (3/8 cells) of cells tested and reduced the increase of AP frequency in the rest of cells (Figure 3B). In contrast, bath application of β-receptor agonist isoproterenol (10-50 µM) did not increase AP firing in all cells tested (n=6, Figure 3C). These results suggest a partial inhibitory effect of β-receptor antagonist. The lack of an agonist effect could indicate saturation of β-receptors activity in the basal condition, while a non-specific action on other sites is also possible. Similar to the β-blocker, the α1 antagonist doxazosin also reduced basal AP rate by 92.8±11.8% and prevented AP induction in 50% of (4/8 cells) neurons (Figure 3C). In contrast to the β-agonist, the α1 agonist (10-50 µM) phenylephrine (PE) partially mimicked the Epi effect and increased spontaneous AP discharge in 66% (6/9 cells) of NA cells (Figure 3D), while without effect in the remainder. These results collectively indicate the predominant roles of α1-receptors in epinephrine-dependent NAm activation pathways; the β-pathway may contribute to the maintenance of basal excitability, but is unlikely to contribute to AP induction.

**Figure 3.**
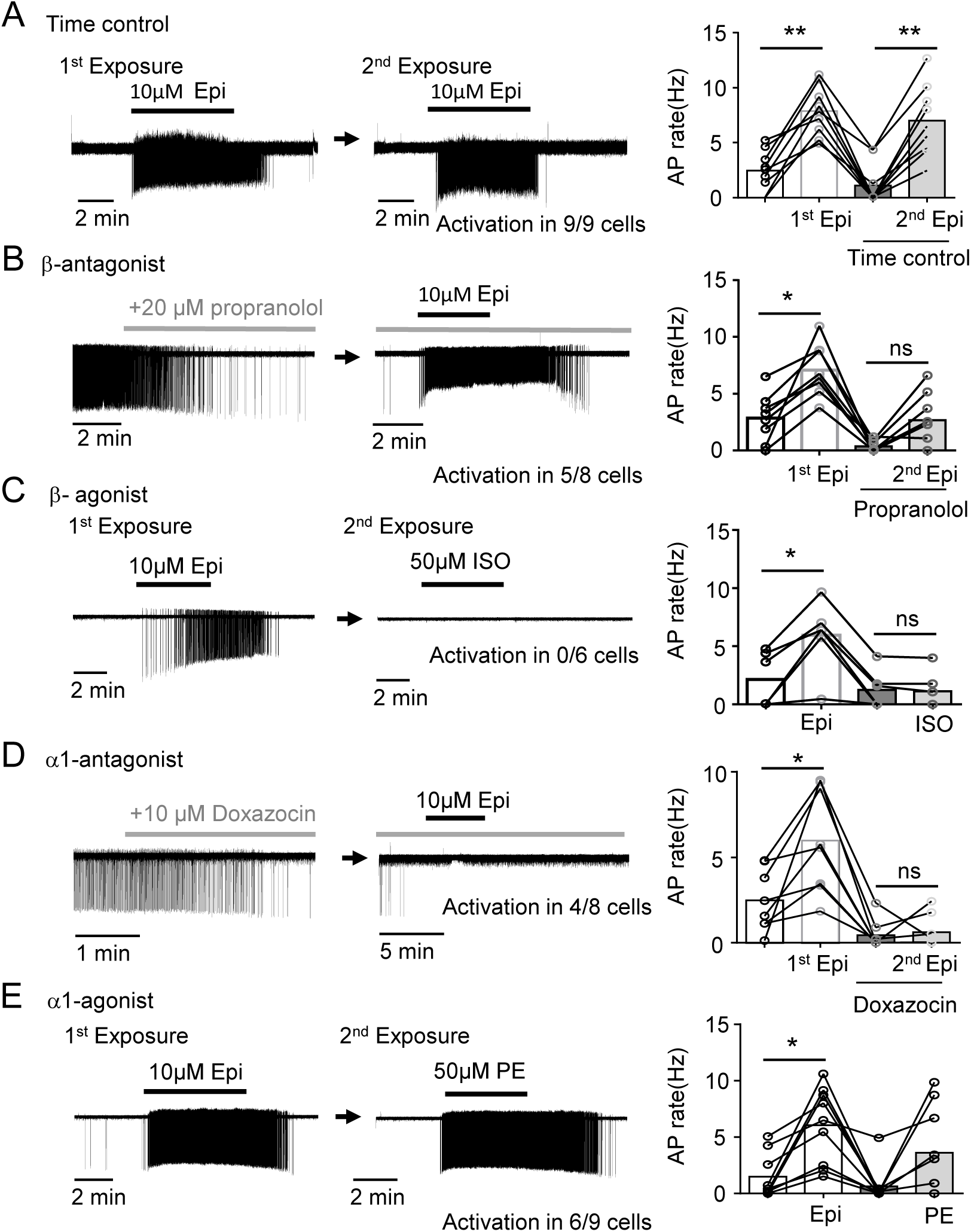
α1 β and adrenergic receptors differentially contribute to NAm excitability. NAm responses to adrenergic agonist and antagonists. Single NAm neurons were exposed to Epi twice, first in control condition and second in the presence of test compound. The AP rate (Hz) indicates the maximum instantaneous AP frequency calculated from multiple 10 second bins. **A.** A time control experiment. Epi exposure (10 µM, 3-5 minutes, shown in horizontal lines) reproducibly activated NAm neurons without rundown of peak firing frequency (repeated-measures ANOVA F(3, 8)=29.63, p<0.0001). **B.** Bath application of β-blocker propranolol reduced basal firing rate and attenuated adrenergic activation (occluded in 3/8 cells, repeated-measures ANOVA F(3, 7)=22.39, p< 0.0001). left: response to propranolol, right: same cell exposed to Epi in the presence of propranolol. **C.** In contrast, β agonist isoproterenol failed (10-50 µM) to induce AP in all cells tested (activation 0/6 cells, repeated-measures ANOVA F(3, 5)=20.64, p= 0.0010). **D.** α1 blocker (Doxazosin) also reduced spontaneous AP rate and attenuated or prevented NAm activation by Epi in 4/8 cells (repeated-measures ANOVA F(3, 7)=19.46, p=0.0002). left: response to doxazosin, right: same cell exposed to Epi in the presence of doxazosin. **E.** Unlike β-agonist, α1 agonist phenylephrine alone could activate 66% (6/9 cells) of NAm neurons (repeated-measures ANOVA F(3, 8)=9.002, p=0.0024). Representative traces (left) and quantitative analysis (right bars) are shown. * p<0.05, ** p<0.01

### AP induction by adrenergic agonists can occur independently of synaptic input

Subtype selective adrenergic agonists/antagonists have been shown to modulate synaptic transmission in NAm cardiac vagal neurons recorded in neonatal rat brainstem slices (Bateman et al. 2012; Boychuk et al. 2011; Philbin et al. 2010; Sharp et al. 2014; Wang et al. 2014). We examined whether such synaptic modulation fully explains the adrenergic activation seen in NAm neurons studied in the young adult mouse.

In a control experiment, direct bath application of 300 µM glutamate reversibly increased AP frequency of NAm neurons (n=2, in agreement with previous studies (Brailoiu et al. 2013a; Brailoiu et al. 2014; Brailoiu et al. 2013c; Sampaio et al. 2014; Yan et al. 2009), supporting the possible contributing role of excitatory synaptic transmission in adrenergic activation of NAm neurons. However this is unlikely to provide the sole explanation. While bath application of NBQX (10 µM), the AMPA/kainate receptor antagonist, significantly decreased pre-drug AP discharge rate in spontaneously active neurons (-58.8±40.4% change from pre-drug baseline frequency, n=6, Figure 4A), Epi still increased the AP firing rate in 83% (5/6 cells) of the NAm neurons (Figure 4A). These results indicate that tonic glutamate receptor activation supports the spontaneous activity of NAm neurons, but is not required for AP induction by adrenergic agonists.

**Figure 4.**
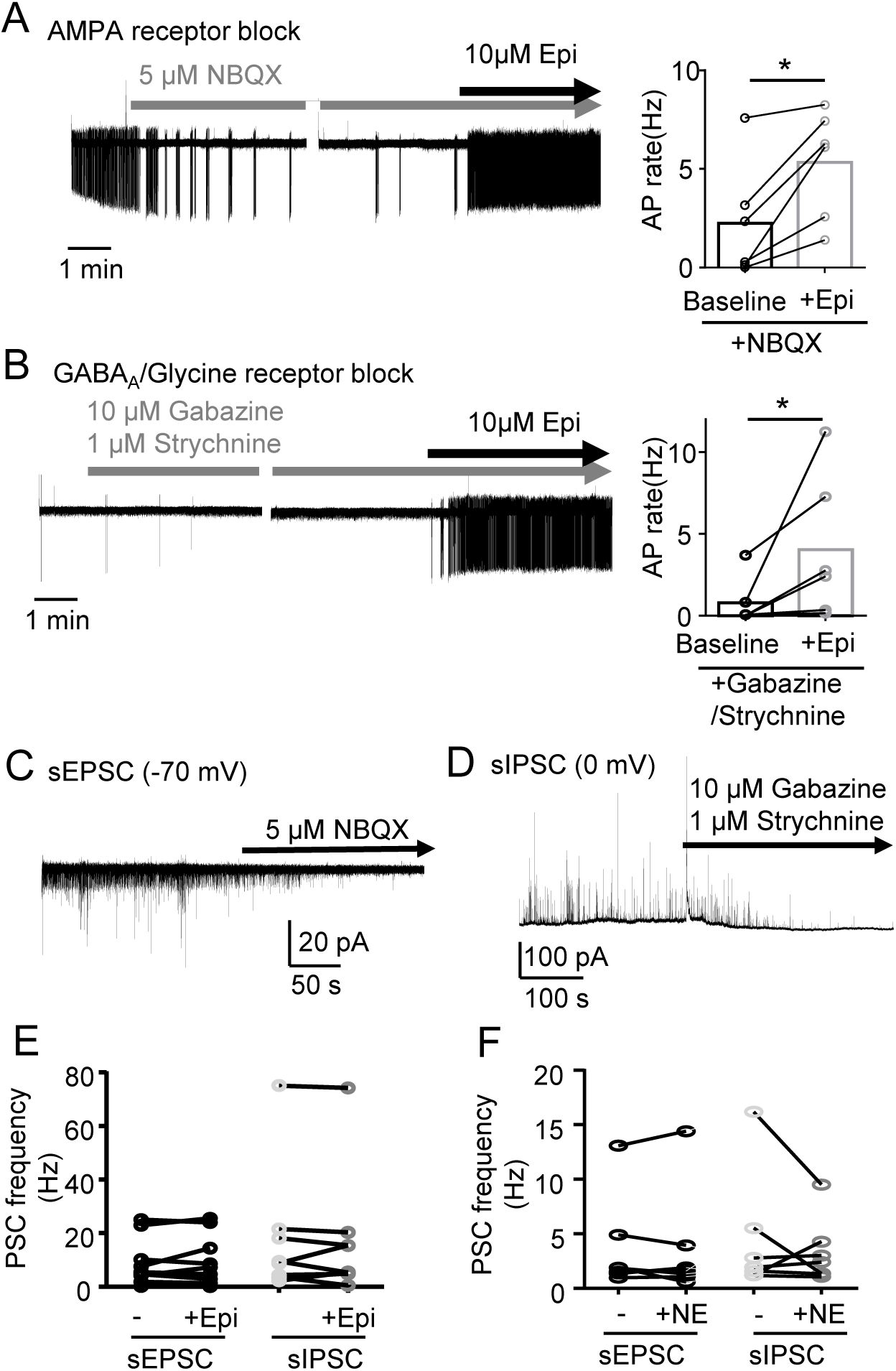
Adrenergic activation does not require synaptic mechanism. **A**. Inhibition of postsynaptic AMPA/kainic acid receptors by NBQX significantly reduced basal firing rate of NAm neurons, but did not prevent activation by Epi (AP rate increase in 5/6 cells, p=0.0313, Wilcoxon matched-pairs signed rank test). Trace: left: response to NBQX, right: same cell exposed to Epi in the presence of NBQX. **B.** Similarly, inhibition of inhibitory synaptic transmission by Gabazine and strychnine did not prevent the adrenergic activation of NAm neurons. (AP rate increase in 6/6 cells, p=0.0313, Wilcoxon matched-pairs signed rank test) Trace: left: response to gabazine and strychnine, right: same cell exposed to Epi in the presence of gabazine and strychnine. **C&D** NBQX and gabazine/strychnine cocktail used in these experiments effectively inhibited spontaneous excitatory (sEPSC) and inhibitory currents (sIPSC), respectively. **E&F** Epinephrine or norepinephrine did not significantly modify sEPSC or sIPSC frequency in NAm neurons under the condition where spontaneous AP was reliably induced (* p<0.05).

We also tested whether inhibitory control was similarly affected, by eliminating inhibitory synaptic currents using co-application of a mixture of the GABA_A_R antagonist gabazine (10 µM) and the Glycine-R antagonist strychnine (1 µM). This synaptic disinhibition did not significantly modulate basal spontaneous AP rate (-11.9±69.5 % change from pre-drug baseline frequency, n=6 cells Figure 4B), and Epi still increased the spontaneous AP rate in all neurons tested (6/6 cells, Figure 4B). These results demonstrate that adrenergic activation of NAm neurons does not rely solely on either excitatory or inhibitory synaptic transmission.

To further examine whether NAm activation by adrenergic agonists might include a synaptic component, postsynaptic currents of the NAm neurons were recorded during Epi or NE exposures. Most of the isolated sEPSC and sIPSC currents were blocked by NBQX (10 µM) and gabazine (10 µM)/ strychnine (1 µM) exposure, respectively (Figure 4C, D). Exposure to 10 µM Epi or NE, which modulated AP firing of the NAm neurons, had no effect on the sEPSC or sIPSC frequency (Figure 4E, F). Thus there was no change in spontaneous synaptic transmission induced by adrenergic agonists in a condition where the spontaneous AP firing rate of NAm neurons was increased.

Together these results suggest that although excitatory synaptic transmission onto NAm neurons can contribute to their spontaneous discharge, it is not critically required for adrenergic activation of NAm neurons, at least in isolated *in vitro* slices.

### Modulation of intrinsic membrane excitability by adrenergic agonists

We next examined whether modulation of intrinsic membrane excitability could be a potential contributor to the adrenergic activation of NAm neurons. We characterized excitability of NAm neurons by whole-cell recordings. Similar to previous studies using preparations from mature animals (Johnson and Getting 1991; 1992; Mendelowitz 1996), depolarizing current evoked a train of regular APs interleaved with a large immediate after-hyperpolarization, likely mediated by SK channels (Lin et al. 2011). A brief hyperpolarizing current typically accompanied by a membrane voltage sag and followed by a rebound AP immediately after termination of hyperpolarizing current was seen in 88% of neurons (88/100 cells, Figure 5A). There was variability in the rebound AP shape during a spike train; one showing an immediate, small afterhyperpolarization (e.g. Figure 5A) can be contrasted with another marked by a larger after-hyperpolarization (e.g. Figure 5B). In both cases, the rebound depolarization was sensitive to intracellular dialysis and became undetectable within ~3 minutes after membrane break-in (Figure 5A). Previous *in situ* hybridization analyses showed expression of polyamine biosynthesis enzymes within the NAm (Figure 5C), thus it is possible that wash-out of endogenous polyamine may contribute to the rundown effect. In fact, the rundown could be partly prevented when 0.5-1 mM spermine was supplied in the intracellular solution (Figure 5B), suggesting that endogenous polyamines or related molecules maintain excitability (see Discussion). Spermine (0.5 mM) was therefore routinely included in subsequent whole-cell current clamp recordings.

**Figure 5.**
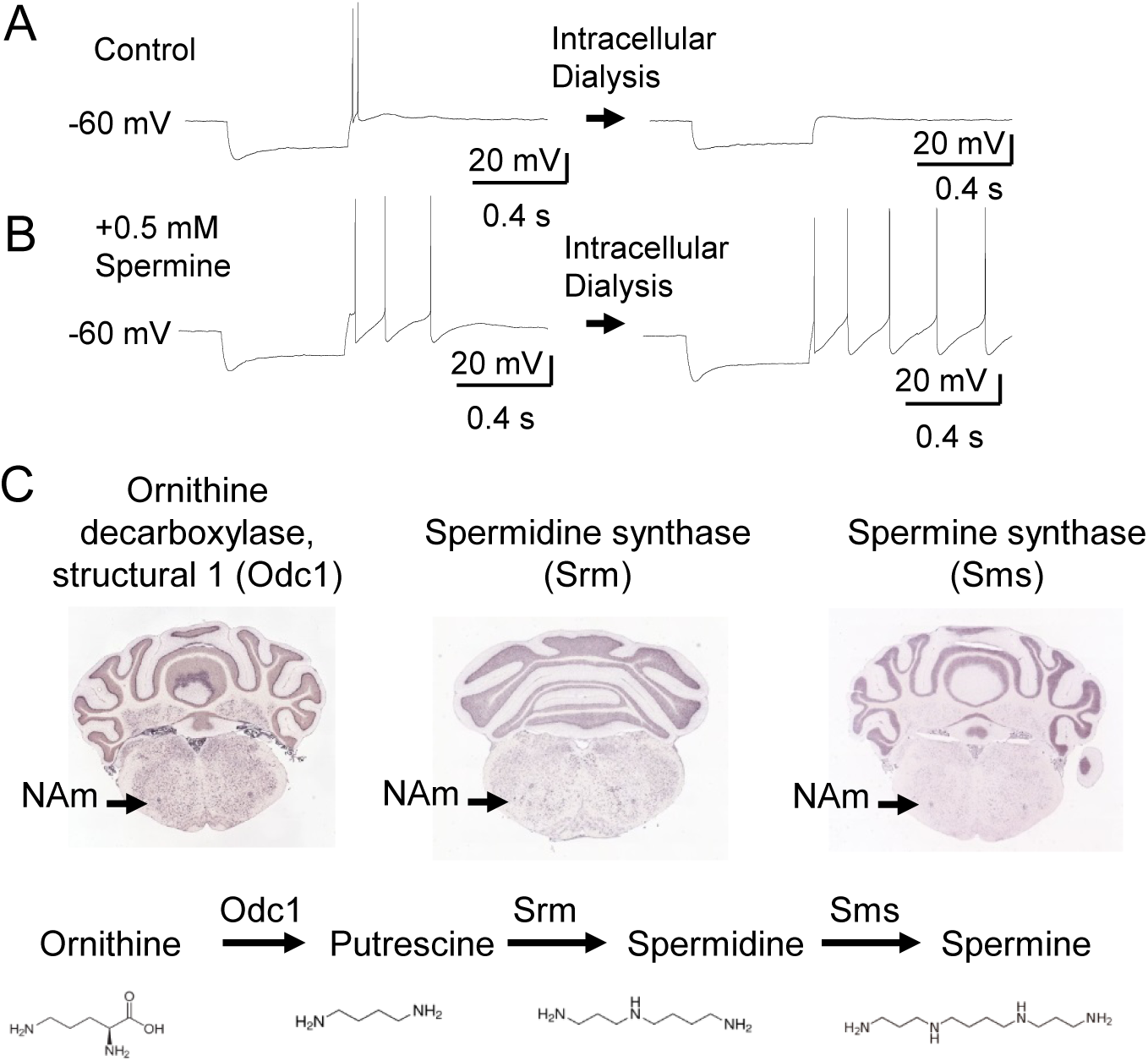
Modulation of intrinsic excitability by adrenergic stimulation. **A&B** Intracellular dialysis sensitivity of NAm neurons. Rebound discharge was sensitive to intracellular dialysis and typically disappeared within 5 minutes after break-in. (**A**). This run-down effect could be reduced by inclusion of 0.5 mM spermine (**B**). **C.** Enzymes involved in polyamine biosynthesis (Odc1, Srm, Sms) are expressed in NAm (data from Allen Brain Atlas).

Bath application of Epi and NE significantly modified intrinsic excitability of NAm neurons (Figure 6A-E). A representative response is shown in Figure 5C. Consistent with extracellular recordings, Epi exposure triggered continuous spontaneous discharges in whole-cell recorded NAm neurons when held at -60 mV. In order to avoid spontaneous AP generation, intrinsic membrane excitability was determined at -70 mV. Membrane resistance was measured by applying small hyperpolarizing voltage pulses, in order to evaluate the amount of leak conductance. As shown in Figure 6B and C, Epi exposure significantly modulated excitability of NAm neurons. Epi and NE increased membrane resistance (Epi: 24.9±11.4% (n=10) and NE: 17.7±16.0% (n=13), while no changes were seen in control experiments (-15.4±21.6%, n=7, Figure 6D). The increased membrane resistance suggests increased excitability by closure of leak currents. This was accompanied by a larger membrane sag during hyperpolarization mediated by Ih current, and enhancement of rebound depolarization which frequently resulted in AP generation mediated by T-type calcium channels, even at -70 mV (see below). The change in rebound AP was robust and a single brief hyperpolarization pulse (<500 ms) could evoke recurrent AP firing lasting 3-5 seconds (see Figure 8C). On the other hand, Epi and NE did not change the numbers of AP evoked by depolarizing current pulses (Figure 6E). Overall, these adrenergic agonists selectively increased some, but not all aspects of intrinsic membrane excitability.

**Figure 6.**
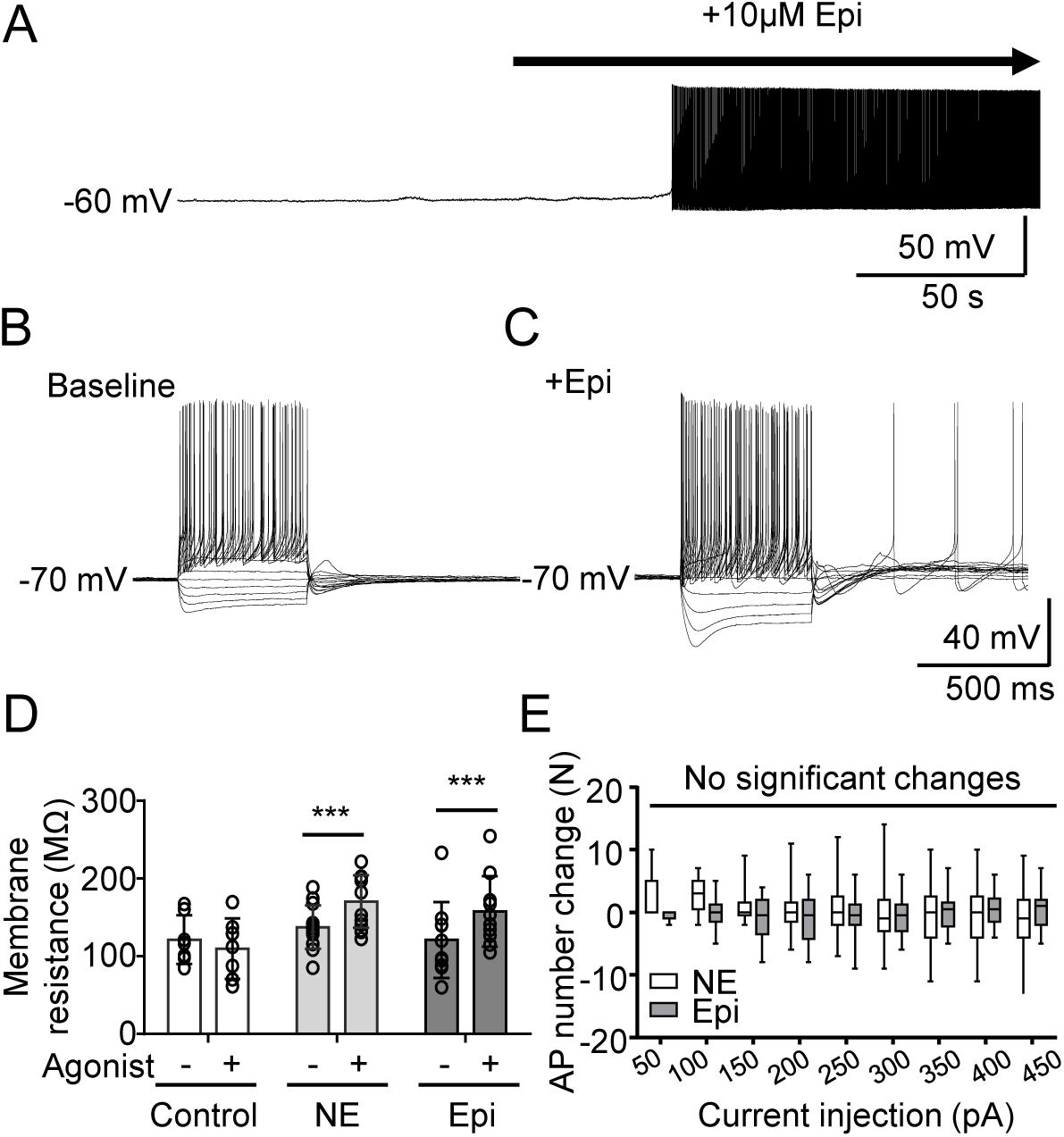
Adrenergic activation of intrinsic membrane excitability of NAm neurons. **A.** adrenergic activation recorded in a whole-cell clamped NAm neuron. Exposure to Epi led to spontaneous AP firing. **B&C.** Voltage responses to step current injection (-200 pA, +50 pA increments) in the same neurons in baseline (**B**) and after exposure to epinephrine (**C**). Note that measurements were made at -60 mV to prevent generation of spontaneous action potentials. **D.** Membrane resistance changes before and after application of each adrenergic agonist. Control n=7. NE n=13, Epi n=10, repeated-measures ANOVA F(1, 44) =36.87, p<0.0001, *** p<0.005. **E.** Changes in evoked action potential numbers after NE or Epi application. No significant changes were detected. NE: n=13, Epi: n=10, repeated-measures ANOVA F(1, 21) = 0.4821, p =0.495.

### Ih-current in NAm neurons

The voltage-sag observed during hyperpolarization (Figure 7A) indicates the presence of hyperpolarization-activated current (Ih) current, and a previous study in guinea pig suggested that cesium-sensitive putative Ih current (Q-current) is involved in rebound depolarization of NAm neurons (Johnson and Getting 1991). Ih is required for spontaneous AP generation in various neurons (Resch et al. 2017), however this was not the case in the NAm. We found that the selective Ih inhibitor ZD7288 (10 µM) completely abolished the voltage sag (n=3, Figure 7A). However, exposure to ZD7288 had no significant effect on the basal firing rate (4.2±74.0%, n=10), and Epi remained effective at inducing AP firing in all NAm neurons tested (n=6 in repetitive experiments Figure 7B&C, and 4 experiments with single Epi exposure (data not shown)). Thus Ih is functionally expressed in NAm cholinergic neurons, but is not required for adrenergic activation of these cells.

**Figure 7.**
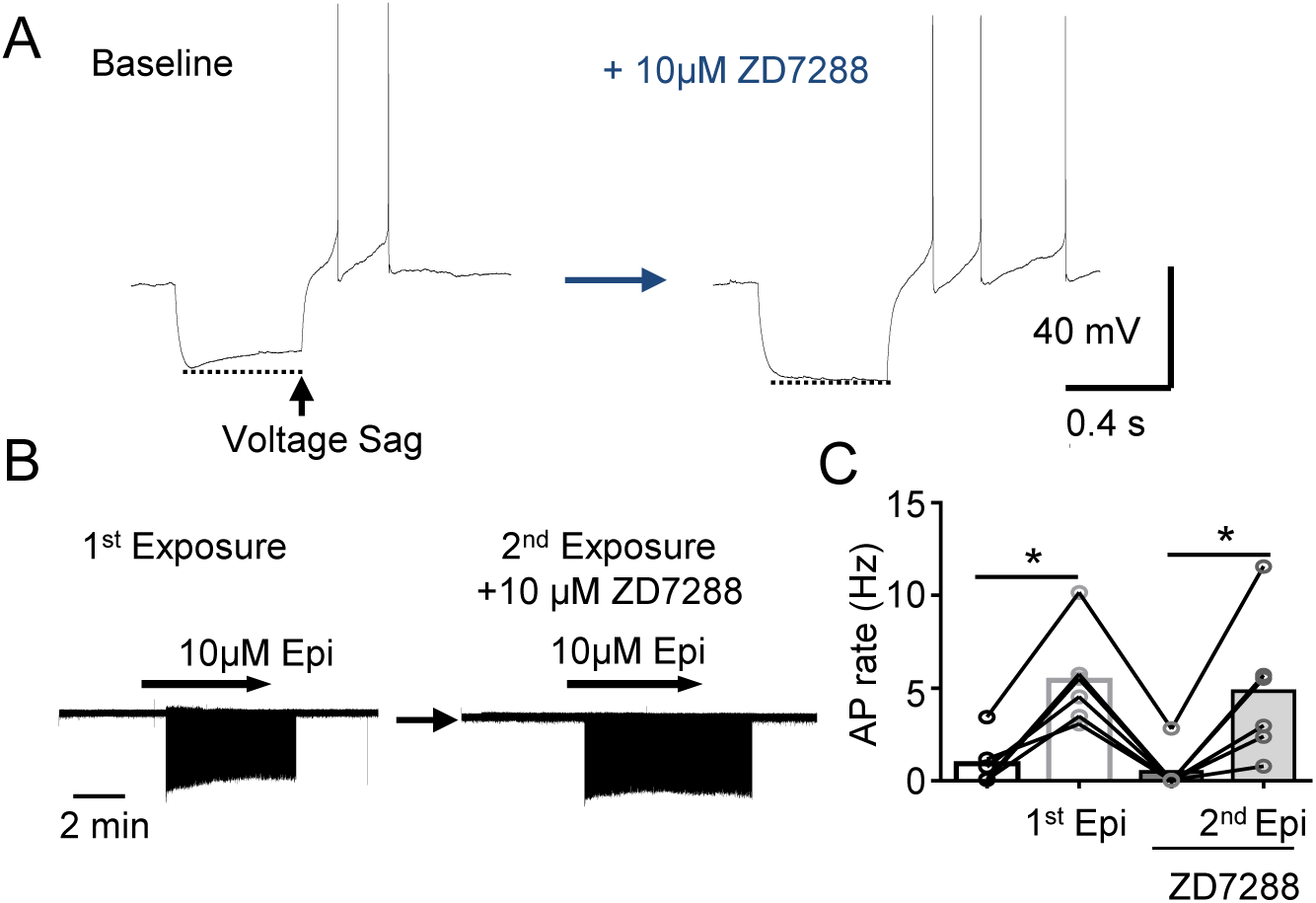
Hyperpolarization activated current (Ih) is expressed but not required for adrenergic activation of NAm neurons. **A.** Ih-mediated membrane sag during hyperpolarizing current was abolished by Ih inhibitor ZD7288 (10 µM). left: control, right: after ZD7288 treatment. **B&C.** ZD7288 at this concentration did not prevent NAm activation by epinephrine **B.** Representative traces. left: exposure to Epi, right: second exposure to Epi in the presence of ZD7288 in the same cell. **C.** Quantitative analysis. (n=6, repeated-measures ANOVA F(3, 5)=17.32, p=0.0020, *p<0.05).

### T-type calcium current in NAm neurons

T-type calcium channels are expressed in various pacemaking cells and contribute to spontaneous AP discharges (Chemin et al. 2002; Perez-Reyes 2003). We next examined the potential role of T-type Ca^2+^ current (ICaT) in NAm excitability. In voltage clamp recordings, low voltage-activated current was reliably evoked by a voltage protocol with a mean current density 4.90±1.28 pA/pF (Figure 8A), a half-activation voltage of - 54.63±1.24 mV, and a half-maximum steady stationary inactivation of -77.56±0.63 mV (Figure 8B). The low voltage-activated current is typically mediated by T-type calcium channels. In fact, T-type calcium channels genes Cav3.1/Cacna1g and Cav3.2/Cacna1h are likely expressed in the NAm (Figure 8C). The selective ICaT inhibitors TTP-A2 (Kraus et al. 2010) and Z944 (Tringham et al. 2012) (1 µM, n=2 for each drug) abolished the inward current as well as the generation of rebound AP’s in both control solution and following Epi exposure (Figure 8D). The ICaT significantly contributes to NAm excitability, since the inhibitor Z944 reduced the spontaneous AP discharge rate by 59.9±33.1% (n=6). However, elimination of ICaT by Z944 did not fully prevent NAm activation by Epi (regular spiking (n=4/7) or burst firing (n=2/7) (Figure 8E), except in one of the regular spiking neurons tested (1/7 cell). Therefore, while ICaT contributes to basal NAm neuronal excitability, it is not required for Epi-dependent NAm neuronal activation.

**Figure 8.**
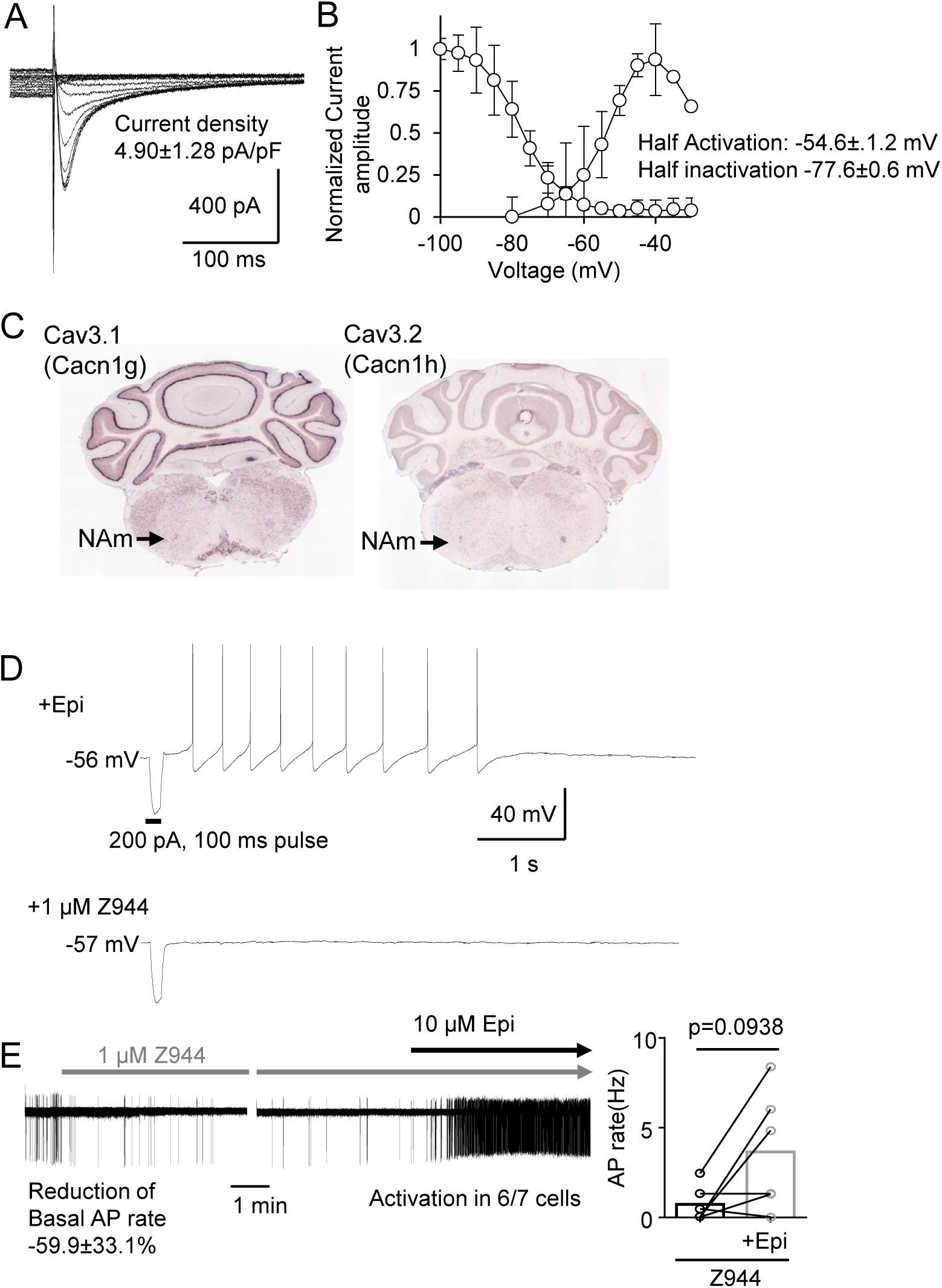
Contribution of T-type Calcium current (ICaT) to the intrinsic membrane excitability of NAm neurons. **A.** Representative traces of the steady current inactivation of Isolated ICaT current of the NAm neuron. **B.** Summary plots of ICaT activation/inactivation kinetics. **C.** Cav3.1 (*Cacna1g*) and Cav3.2 (*Cacna1h*) mRNAs are detected in the NAm (data from Allen Brain Institute). **D** ICaT current inhibitor Z944 (1 µM) abolished rebound action potentials of NAm neurons. **E.** ICaT inhibition significantly reduced basal AP firing rate of NAm neurons, but did not fully prevent NAm activation by epinephrine (AP rate increase in 6/7 cells, p=0.094, Wilcoxon matched-pairs signed rank test). Left trace shows effect of Z944 on basal AP firing and induction of AP by Epi in the same recorded neuron. Quantitative analysis is shown in bar graph.

### Persistent sodium current in NAm neurons

We analyzed the persistent sodium current (INaP) in adrenergic activation of Nam neurons. INaP is a slow inward current carried by voltage-gated sodium channels, and contributes to spontaneous discharges in various cell types (Bevan and Wilson 1999; Khaliq and Bean 2010, see Discussion; Koizumi and Smith 2008). INaP was isolated as a 1 µM TTX sensitive component of inward current that developed during a slow voltage ramp (+20 mV/s, 33.6±24.1 pA at -40 mV, n=5 Figure 9A). The isolated INaP emerged near the resting membrane potential of NAm neurons (Figure 9A), and continuously developed to peak at -10~0 mV. The non-selective persistent sodium current inhibitor riluzole (30 µM, IC_50_ 2.3~51 µM (Wang et al. 2008; Zona et al. 1998)) was highly effective at preventing adrenergic activation of NAm neurons and fully prevented the activation of AP discharges in all 8 NAm neurons tested (Figure 9B). It is notable that even in the presence of riluzole, AP’s could be evoked by local electrical stimulation, indicating that the observed effect is not due to a non-selective inhibition of sodium current. We further evaluated the contribution of INaP using a specific INaP inhibitor GS967 (Belardinelli et al. 2013) at the concentration of 1 µM where the drug has high selectivity for INaP while may not fully inhibiting the current in these neurons (Anderson et al. 2014; Belardinelli et al. 2013). INaP inhibition by GS967 fully prevented AP induction in 3/6 cells tested and severely attenuated the response to Epi in the rest of cells (Figure 9C). These results suggest an important contribution of INaP current to the adrenergic activation of NAm neurons.

**Figure 9.**
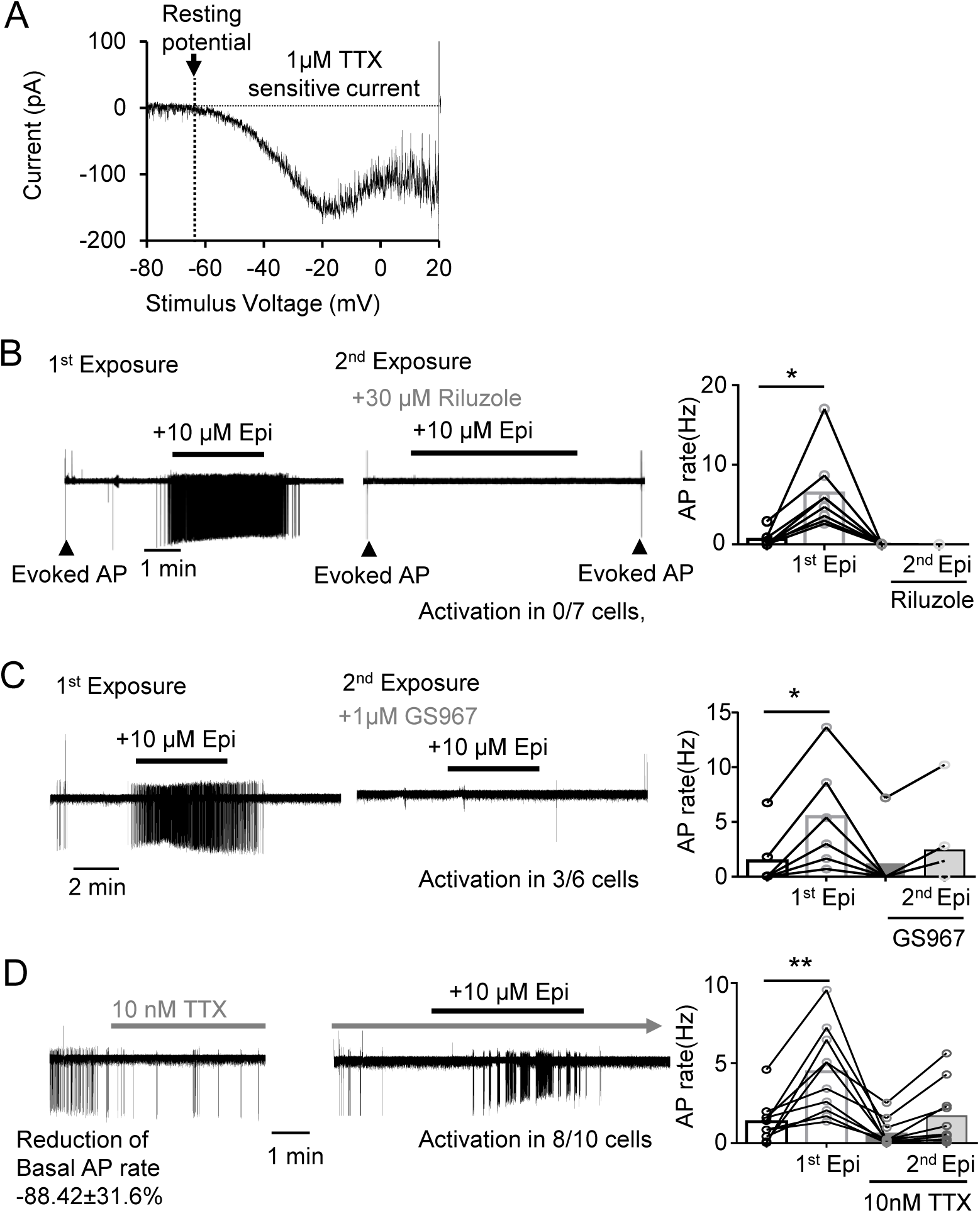
Persistent sodium current (INaP) is required for the adrenergic activation of NAm neurons. **A.** INaP in a NAm neuron isolated as TTX sensitive inward current generated during a slow voltage ramp (20 mV/s). B-C. Effect of sodium channel blocker on the AP induction by Epi. **B.** Non-selective INaP inhibitor riluzole effectively prevented NAm activation by epinephrine (activation in 0/8 cells, repeated-measures ANOVA F(3, 7)=14.23, p=0.0057). Note that riluzole did not prevent action potentials (AP) evoked by electrical stimulation of surrounding tissue (arrow heads). left: exposure to Epi, right: second exposure to Epi in the presence of riluzole in the same cell. **C.** A low concentration of INaP selective inhibitor GS967inhibited activation by epinephrine in 50% (3/6 cells) and reduced it in the remaining of cells. repeated-measures ANOVA F (3, 5) = 24.76, p<0.0001. left: exposure to Epi, right: second exposure to Epi in the presence of GS967 in the same cell. **D.** A low concentration of TTX (10 nM) reduced spontaneous AP, while Epi still increased AP firing rate in the 80% (8/10 cells) of NAm cells tested. repeated-measures ANOVA F(3, 9)=11.25, p=0.0009 Representative traces (**left**) shows effect of TTX on basal firing and response to Epi in the presence of TTX in the same cell. Quantitative analysis is shown in bar graph (**right**). * p<0.05, ** p<0.01

We next examined the contribution of TTX insensitive sodium currents to the adrenergic activation. Bath application of 10 nM TTX severely lowered the spontaneous AP rate of NAm neurons (88.4±31.6% reduction from baseline, n=10, Figure 9D), indicating a strong contribution of TTX-sensitive Na^+^ current to basal excitability. However, Epi was still able to increase AP firing rate in 80% of NAm neurons (8/10) tested under this condition (Figure 9D). Thus, while sodium channels that are highly sensitive to TTX are important for basal NAm excitability, TTX-insensitive sodium channels alone are sufficient to mediate Epi-dependent AP increases in NAm neurons.

Finally, we examined the effectiveness of the non-selective sodium current inhibitor flecainide, an antiarrhythmic useful in the control of a variety of supraventricular cardiac tachyarrhythmias, including those bearing *SCN5A* mutations underlying Brugada Syndrome (Salvage et al. 2017). Flecainide (20 µM) was partially effective at preventing AP induction, and blocked Epi activation in 3/5 cells tested. Thus in addition to multiple myocardial targets, including Nav1.5 channels in the septal conduction system and cardiac myocytes, this clinically useful peripheral sodium channel blocker may also reduce central vagal nerve activity.

Together these results suggest that adrenergic activation of NAm neurons requires a persistent sodium current, predominately mediated by sodium channels with low TTX-sensitivity.

## Discussion

This study identifies a central adrenergic mechanism regulating the outflow of preganglionic cardiac vagal neurons in the brainstem. We found that two adrenergic agonists, epinephrine and norepinephrine, significantly increase the spontaneous discharge rate of these neurons by acting at α1 β and receptors (Figures 2&3). This adrenergic activation, however, does not require synaptic transmission, since 1) it occurred even when the postsynaptic receptors were pharmacologically inhibited, and 2) following agonist exposure, there was no significant change in the frequency of transmitter release at either excitatory and inhibitory synapses onto NAm neurons (Figure 4). Instead, we observed an increase in their intrinsic membrane excitability leading to vigorous rhythmic discharge (Figures 6). We also found that mouse NAm neurons express functional hyperpolarization-activated current (Ih), T-type calcium current (ICaT), and persistent sodium (INaP) current, and of these, the INaP is an important contributor to the intrinsic adrenergic activation, while changes in the other ionic currents are likely involved. These results define a novel mechanism for central adrenergic regulation of parasympathetic output to the heart, and define a target excitatory mechanism underlying NAm preganglionic neuron rhythmic activity.

The origin of the adrenergic input remains under study. Because of poor blood brain barrier permeability of circulating catecholamines (Kostrzewa 2007), brainstem adrenergic neurons located in proximity to NAm (e.g. A1, C1 groups) are a more likely source of synaptic regulation. Optogenetic activation of neurons within the C1 group in the medulla triggers mild bradycardia (Abbott et al. 2013), and the adrenergic mechanism identified in this study may contribute in part to this effect. Given the major role of local brainstem adrenergic neurons in sympathetic cardiac regulation, the adrenergic NAm activation described in this study may be a collateral pathway to counterbalance excessive cardiac sympathetic outflow to limit tachycardia. Alternatively, abnormal co-activation of the cardiac sympathetic and parasympathetic pathways might increase the risk of arrhythmias such as atrial fibrillation (Inoue and Zipes 1987; Ogawa et al. 2007).

Previous studies have demonstrated that adrenergic signaling suppresses inhibitory transmission to cardiac vagal neurons. Thus β-receptor agonists inhibit both sEPSCs and sIPSCs (Bateman et al. 2012), while α1-agonists (Boychuk et al. 2011) and α2-agonists inhibit sIPSC (Philbin et al. 2010; Sharp et al. 2014). Finally, optogenetic activation of adrenergic neurons within the locus coeruleus inhibits NAm neuronal discharge by selective augmentation of inhibitory currents (Wang et al. 2014). Based on these reports, it was unexpected that exposure to adrenergic agonists had no significant effect on afferent NAm synaptic transmission in our preparation (Figure 4). The discrepancy could be explained by the 1) species differences (rat vs mouse), 2) the age of animals (neonate/juvenile vs young adult), and 3) the precise neuronal population (retrograde tracer labeling vs Chat-cre reporter) recorded. Among these, age may play a significant role, since rodents show significant postnatal development of the parasympathetic autonomic regulatory system (Kasparov and Paton 1997) during the initial 3-weeks of life. In addition, we used thinner brain slices (200 µm) from the larger adult brainstem than previous studies (500-600 µm) using the smaller brainstem of younger animals where neuronal and synaptic density could be higher, leading to alternative estimates of the contribution of synaptic input.

In addition, our studies reliably detected rebound APs (Figure 5) which were not reported in studies using preparations from immature animals (Mendelowitz 1996; Mihalevich et al. 1996). Since rebound APs have been observed in adult guinea pig NAm neurons (Johnson and Getting 1991), there could be age-dependent development of this intrinsic membrane excitation mechanism. Alternatively, membrane loading with the lipophilic dye used for cardiac retrograde tracing used in previous studies (Mendelowitz 1996; Mihalevich et al. 1996) may have modified excitability of the plasma membrane and masked the intrinsic membrane excitatory mechanism.

A previous study of NAm neurons in adult guinea pig brainstem, using sharp electrode intracellular recording (Johnson and Getting 1991), also identified multiple neuronal populations according to membrane excitability and cell morphology. The pool of preganglionic neurons recorded in this study is consistent with one of the populations carrying a delayed rebound action potentials, and the other population may constitute the smaller Chat-tdtomato^+^ non-preganglionic neurons and Chat-tdtomato^-^ neurons (Figure 1). Our results imply cardiac preganglionic NAm neurons can be further subdivided based on their response to adrenergic agonists; neurons that show a simple increase in regular spiking and those that generate a burst discharge (Figure 2). Because of the low chance of encountering them, we could not characterize the burst firing neurons in further detail. Thus it is still unclear whether these two responses may represent two different intrinsic cell populations or a difference attributable to synaptic connectivity. However since changes in intrinsic membrane excitability were consistently observed in all neurons tested (Figure 5), it is likely that enhancement of membrane excitability by adrenergic agonists affects both cell types.

Our study suggests that α1-receptors are the major mediator of the adrenergic NAm excitability increase, while β-receptors may be important for maintenance of basal membrane excitability. The contribution of these receptors to neuronal membrane excitability has been demonstrated in other neurons (Grzelka et al. 2017; Martinez-Pena y Valenzuela et al. 2004; Randle et al. 1986). The α1-and β-adrenoceptors are typically coupled with distinct GPCR signaling (i.e. α1-Rs with Gq, β-Rs with Gs). However these GPCR pathways crosstalk with each other, and β-receptor activation can activate Gq signaling pathway by direct (Wenzel-Seifert and Seifert 2000; Zhu et al. 1994) or indirect mechanisms (Cervantes et al. 2010; Galaz-Montoya et al. 2017), which may in part explain how distinct adrenoceptors similarly contribute to NAm intrinsic excitability.

Our study also identified multiple ionic currents involved in oscillatory membrane discharges in NAm neurons. We confirmed functional ICaT expression in the NAm and its contribution to rebound discharge generation. Consistent with electrophysiological detection, NAm neurons show high expression of Cav3.1 and Cav3.2 mRNA transcripts (Figure 8C, Allen Brain Atlas). In addition to ICaT, NAm neurons express a significant Ih current which is involved in membrane oscillation of some cells. While these currents did not appear to contribute to adrenergic activation in the *in vitro* preparation, these properties could be involved in complex physiological processes such as vagal outflow during baroreflex and sustained vagal activity *in vivo*.

Our pharmacological studies (Figure 9) suggest that INaP is an important component of adrenergic activation of NAm neurons. INaP, similar to the one we identified here, contributes to the pacemaking discharge of many neurons (Beurrier et al. 1999; Bevan and Wilson 1999; Khaliq and Bean 2010; Koizumi and Smith 2008; Mercer et al. 2007). INaP is also a mediator of muscarinic agonist-induced spontaneous AP generation of hippocampal CA1 neurons (Yamada-Hanff and Bean 2013), and NAm neurons also possess a similar INaP dependent metabotropic activation mechanism.

While the specific sodium channels responsible for INaP are not yet determined, our results suggest that TTX-insensitive sodium channels alone are sufficient to mediate adrenergic activation of NAm (Figure 9). TTX-insensitive channels are encoded by Scn5a (Nav1.5), Scn10a (Nav1.8), and Scn11a (Nav1.9) (Goldin 2001) and are predominantly expressed in cardiac myocytes and peripheral nerves. Brainstem expression of these sodium channels likely contribute to the unique excitation mechanism of NAm, and may be fortuitously affected by peripheral sodium channel blockers, such as flecainide.

In summary, our study shows an adrenergic activation mechanism of NAm neurons mediated by modulation of intrinsic membrane excitability that requires INaP current. The results add an additional site of interaction between the sympathetic and parasympathetic systems within the medulla (DePuy et al. 2013), mediating the finely tuned balance of central autonomic cardiovascular regulation.

## Acknowledgments

We thank Dr Terry Snutch at the University of British Columbia for generously providing Z944.

## Grants

AHA 14POST20130031(IA), NIH/NINDS U01 NS090340 (JLN)

